# Commensal-derived short-chain fatty acids disrupt lipid membrane homeostasis in *Staphylococcus aureus*

**DOI:** 10.1101/2024.08.12.607382

**Authors:** Joshua R. Fletcher, Lisa A. Hansen, Richard Martinez, Christian D. Freeman, Niall Thorns, Alex R. Villareal, Mitchell R. Penningroth, Grace A. Vogt, Matthew Tyler, Kelly M. Hines, Ryan C. Hunter

**Author notes:** To whom correspondence should be addressed: Ryan C. Hunter Department of Microbiology & Immunology Jacobs School of Medicine and Biomedical Sciences University at Buffalo 955 Main St. Room 5120 Buffalo, NY 14203.

## Abstract

The role of commensal anaerobic bacteria in chronic respiratory infections is unclear, yet they can exist in abundances comparable to canonical pathogens *in vivo*. Their contributions to the metabolic landscape of the host environment may influence pathogen behavior by competing for nutrients and creating inhospitable conditions via toxic metabolites. Here, we reveal a mechanism by which the anaerobe-derived short chain fatty acids (SCFAs) propionate and butyrate negatively affect *Staphylococcus aureus* physiology by disrupting branched chain fatty acid (BCFA) metabolism. In turn, BCFA impairment results in impaired growth, diminished expression of the agr quorum sensing system, as well as increased sensitivity to membrane-targeting antimicrobials. Altered BCFA metabolism also reduces *S. aureus* fitness in competition with *Pseudomonas aeruginosa*, suggesting that airway microbiome composition and the metabolites they produce and exchange directly impact pathogen succession over time. The pleiotropic effects of these SCFAs on *S. aureus* fitness and their ubiquity as metabolites in animals also suggests that they may be effective as sensitizers to traditional antimicrobial agents when used in combination.

---

*Staphylococcus aureus* is a Gram positive pathogen often found in polymicrobial infections of the cystic fibrosis (CF) lung and upper airways of individuals with chronic rhinosinusitis (CRS)(*1–5*). Despite its arsenal of virulence factors and association with respiratory infections, *S. aureus* is also commonly present in the airways of healthy individuals to no obvious detriment(*6, 7*). Since *S. aureus* plays a dual role as both pathogen and commensal, it is key to understand how it integrates environmental cues to regulate its metabolism and virulence.

A common feature of both CF and CRS is the accumulation of airway mucus, resulting in hypoxic microenvironments and colonization by anaerobic bacteria that can utilize mucin glycoproteins as growth substrates(*5, 8, 9*). As a result, anaerobes generate short chain fatty acids (SCFAs) that stimulate host inflammation, serve as carbon sources for pathogens like *Pseudomonas aeruginosa*, or in the case of propionate and butyrate, impair *S. aureus* growth(*8, 10–13*). Mechanisms underlying SCFA-mediated inhibition of *S. aureus* are unknown, but evidence from our group and others implicates lipid metabolism and cell wall stress(*11–13*). For instance, perturbation of teichoic acids renders *S. aureus* susceptible to inhibition by propionate *in vitro* and in a murine wound model(*11*). Moreover, FadX (putatively involved in fatty acid degradation) is required for *S. aureus* growth in propionate, while a *codY* mutant grows significantly better than wildtype in butyrate(*13*). CodY is a master regulator of metabolism and virulence, and among the most highly expressed genes in a *codY* mutant are involved in branched chain amino acid (BCAA) biosynthesis(*14, 15*). BCAAs (isoleucine, leucine, valine) are substrates for branched chain fatty acid (BCFA) production, which are highly abundant in the *S. aureus* membrane; the branched to straight chain fatty acid ratio is essential for regulating membrane fluidity as environmental conditions change(*16–20*). Recently, a *codY* mutant was shown to have elevated anteiso BCFAs in its membrane, and the activity of the Sae two-component system was sensitive to their presence(*21*). Disruption of BCFA production via mutation of branched chain keto acid dehydrogenase (Bkd) results in poor growth, increased membrane rigidity, and sensitivity to environmental stresses(*17–19*).

Given these observations, we hypothesized that propionate and butyrate affect *S. aureus* lipid membrane homeostasis by decreasing BCFA abundance. We found that isoleucine improved *S. aureus* growth in the presence of propionate and butyrate, while leucine and valine had the opposite effect. Mutants incapable of converting isoleucine to anteiso BCFAs exhibited increased sensitivity to both propionate and butyrate, and exogenous isoleucine failed to rescue growth. Consistent with this, targeted lipidomics revealed that *S. aureus* grown in propionate- and butyrate-supplemented media had a lower BCFA to straight chain FA ratio than when grown in LB alone. As a result, SCFAs potentiate the activity of membrane-targeting antibiotics against the type strain (JE2), disrupt quorum signaling, and diminish the competitive fitness of *S. aureus* in co-culture with *P. aeruginosa*. Finally, several *S. aureus* clinical isolates behaved similarly to JE2 across phenotypic assays, indicating that SCFAs act on conserved molecular targets. These findings suggest that during chronic airway infection, commensal-derived SCFAs may synergize with antimicrobials and reduce the competitive fitness of *S. aureus* by impairing BCFA homeostasis.

## Results

### Propionate and butyrate alter the *S. aureus* transcriptome and proteome

Airway microbiota consist of complex communities with several taxa that originate from the oral cavity(*22–24*). This is supported by the relative abundance of *Streptococcus*, *Prevotella*, and *Fusobacterium* spp. in CF sputum and sinus mucus, however, contributions of these strict and facultative anaerobes to airway disease remain poorly understood(*3–5, 25*). In prior work, we found that culturing *S. aureus* in *F. nucleatum* supernatant led to impaired growth that was attributable to the SCFAs propionate and butyrate(*13*). Here, our goal was to further dissect the effects of commensal-derived SCFAs on *S. aureus* to identify mechanisms of action.

We first used a NanoString codeset of 33 probes targeting transcriptional regulators, virulence factors, and metabolic genes to quantify their expression during *S. aureus* JE2 growth in LB supplemented with propionate or butyrate (**Fig. 1a, Suppl. Dataset 1**). Each condition resulted in unique expression patterns, with ten (propionate) and nine (butyrate) transcripts reaching significance versus LB-grown controls (p-adj <0.05). Four transcripts decreased in propionate, while all nine in butyrate increased. Several were previously identified in *S. aureus* grown in *F. nucleatum* supernatants(*13*), despite using a different base medium, underscoring the specific effect of SCFAs on *S. aureus* physiology. Although gene expression in propionate and butyrate was distinct, commonalities were observed between media. For instance, expression of *alsS*, *nanA*, *gltB*, and *fadX* increased in both SCFAs relative to LB alone. *alsS* encodes an acetolactate synthase that produces acetolactate from pyruvate and contributes to isoleucine biosynthesis<u>42</u>. *nanA*, *gltB*, and *fadX* are involved in sialic acid, glutamate, and fatty acidetabolism, respectively(*26, 27*). Additionally, expression of *ilvE*, encoding a CodY-regulated BCAA aminotransferase involved in anteiso BCFA synthesis(*28*)**(Fig. 2a)**, increased in both SCFA media, though it only reached significance in propionate. As BCFAs are abundant in the *S. aureus* membrane, we reasoned that the magnitude of change for transcripts linked to BCFA metabolism may not be as large due to their importance to cellular homeostasis(*18, 21*).

**Fig. 1.**
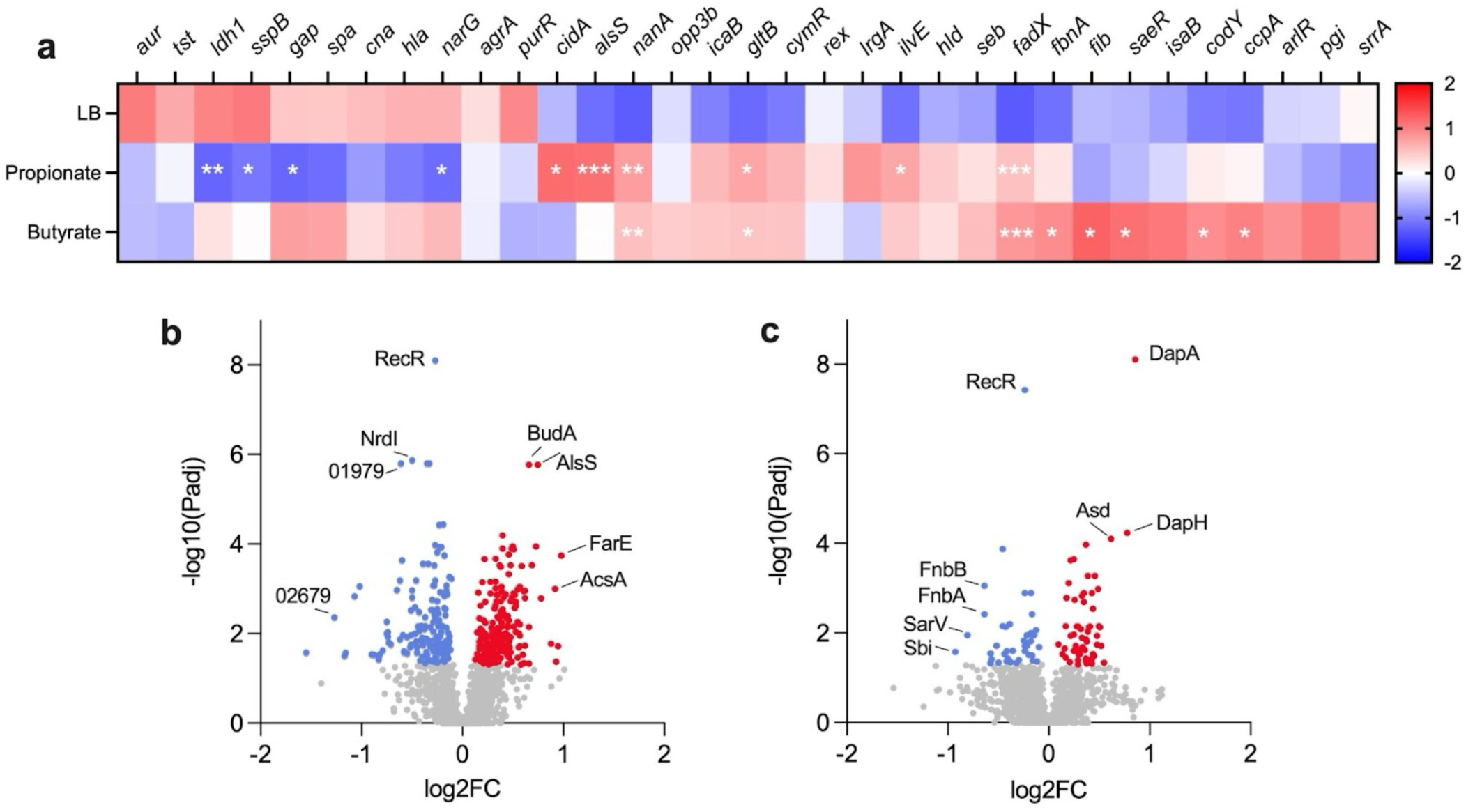
Propionate and butyrate alter *S. aureus* transcript and protein abundances. **(a)** Heatmap depicting Z-scores of log10 transformed NanoString counts from *S. aureus* JE2 grown to OD600 of ∼0.2-0.3 in LB with or without 100 mM of sodium propionate or sodium butyrate (n=3 per condition). Transcripts that exhibited greater than two-fold change in abundance relative to growth in LB were considered statistically significant at a Benjamini-Hochberg adjusted p-value ≤ 0.05 (*** <0.001, ** <0.01, * <0.05). **(b and c)** Volcano plots of the *S. aureus* proteome in LB supplemented with propionate (b) or butyrate (c) compared to LB alone. NanoString and proteomics output can be found in Supplemental Files 1 and 2, respectively.

**Fig. 2.**
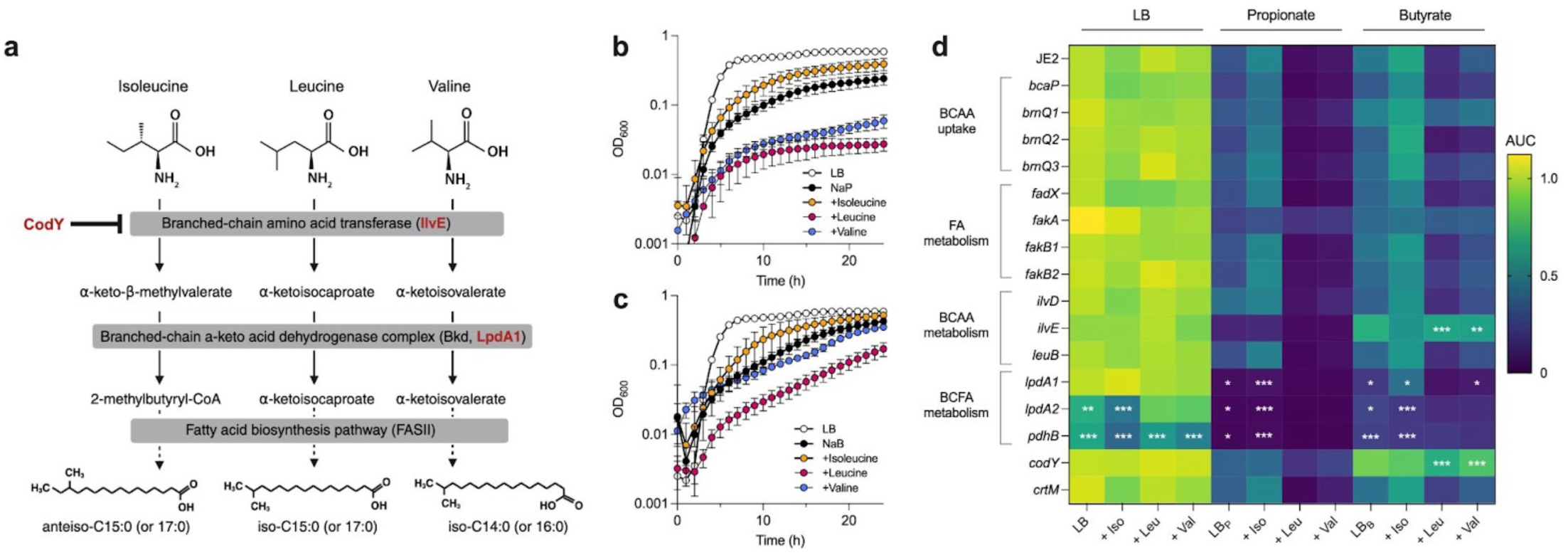
Isoleucine supplementation partially relieves growth inhibition by propionate and butyrate, while leucine and valine enhance it. **(a)** Graphical depiction of the conversion of BCAAs to alpha-keto acids by IlvE, then to CoA-esters by the branched chain ketoacid dehydrogenase (BKD) for their subsequent use as substrates for FASII. Adapted from Chan and Wiedmann (2009). The BKD complex includes LpdA1. **(b, c)** Growth curves of *S. aureus* JE2 in LB with 100 mM of **(b)** sodium propionate (NaP) or **(c)** sodium butyrate (NaB), supplemented with 1 mg/mL of the indicated branched chain amino acid. Error bars represent standard error of the mean for each time point. **d)** Heatmap depicting the normalized area under the curve (AUC) of *S. aureus* JE2 and transposon mutants in genes associated with BCAA uptake/metabolism or fatty acid metabolism (n=4 growth curves per strain, per condition). Data were normalized to JE2 grown in LB alone and are presented as % area under the curve. Large normalized AUC indicates robust growth and is depicted in lighter green to yellow, while low values indicate poor growth and are shown in darker green to dark blue. Statistical significance of each mutant compared to JE2 under a given growth condition was tested using an ordinary two-way ANOVA with Dunn’s multiple comparisons test (*, p<0.05; **, p<0.01; ***, p<0.001). Comparisons were too numerous to depict graphically and are shown in Suppl. File 3.

We next profiled the *S. aureus* proteome under identical conditions (**Fig. 1b,c, Suppl. Dataset 2**). Compared to growth in LB alone, 370 (propionate) and 103 (butyrate) proteins were differentially abundant (p<0.05) with 80 shared between media. RecR was significantly lower when either SCFA was present, as was the NrdI ribonucleotide reductase stimulatory protein. Acetolactate synthase proteins BudA and AlsS were highly induced by propionate only. Conversely, the dihydropicolinate synthases DapA, DapB, and DapH were significantly higher in butyrate but were unaffected by propionate. N-acetylneuraminate lyase (NanA) was induced by propionate, consistent with its transcript levels. In contrast, FbnA was lower in both SCFAs compared to LB alone, despite showing significant transcriptional induction in butyrate. This suggests a disconnect between *fbnA* transcription and translation. Likeise, CodY expression was consistent across media, despite *codY* transcripts being modestly induced by both SCFAs. IlvE was approximately two-fold higher in both SCFA media but was not statistically significant. These transcriptomic and proteomic data indicate that SCFAs have wide-ranging effects on *S. aureus*, with implications for cell envelope homeostasis, metabolism, and pathogenesis. Given our previous work demonstrating a link between SCFAs and lipid metabolism(*13*), we narrowed our focus to BCAAs and BCFAs.

### Isoleucine relieves SCFA-mediated growth inhibition of *S. aureus*

Given (i) increased expression of *ilvE* during growth in SCFAs (**Fig. 1a**), (ii) robust growth of a *codY* mutant in butyrate(*13*), and (iii) increased anteiso BCFAs in the *codY* mutant membrane(*21*), we hypothesized that exogenous BCAAs could mitigate SCFA-mediated growth inhibition by serving as BCFA precursors. To test this, wildtype JE2 was grown in LB supplemented with one BCAA (isoleucine, leucine, or valine) in the presence of propionate or butyrate (**Fig. 2b,c**). Growth was impaired by either SCFA alone, but partially restored by isoleucine. Interestingly, growth was further impaired by leucine and valine, suggesting that iso-BCFAs produced from these substrates are deleterious when SCFAs are present. These data align with previous studies showing that 2-methylbutyric acid (produced by deamination of isoleucine by IlvE) is the preferred substrate of branched-chain ketoacid dehydrogenase (Bkd, **Fig. 2a**), while leucine- and valine-derived intermediates are less efficiently utilized(*20*).

To further investigate these phenotypes, we screened *S. aureus* transposon mutants for propionate and butyrate sensitivity, with and without excess BCAAs. Mutants were selected for their gene product’s involvement in BCAA uptake (*bcaP*, *brnQ1-3*), fatty acid metabolism (*fakA*, *fakB1*, *fakB2*, *fadX*), BCAA metabolism (*ilvD*, *ilvE*, *leuB*), or BCFA metabolism (*lpdA1*, *lpdA2*, *pdhB*). *lpdA2* is downstream of *pdhB* and has not been directly linked to BCFA metabolism. A *codY* mutant was also included given its robust growth in butyrate relative to wildtype(*13*). Finally, we included a staphyloxanthin mutant (*crtM*), as staphyloxanthin can influence membrane fluidity, which is diminished by reduced BCFA content(*16, 17, 29, 30*).

While several mutants displayed altered growth patterns relative to JE2, only *codY*::tn, *ilvE*::tn, *lpdA1*::tn, *lpdA2*::tn, and *pdhB*::tn reached significance in one or more media (**Fig. 2D, Suppl. File 3**). Despite the role of staphyloxanthin in regulating membrane fluidity, *crtM*::tn did not exhibit altered SCFA sensitivity. As expected, *codY*::tn grew significantly better than JE2 in butyrate. *fakA*::tn was less sensitive than JE2 to the additive effect of leucine or valine on growth impairment by propionate and butyrate, while *brnQ1*::tn exhibited this insensitivity in butyrate only. Interestingly, *ilvE*::tn phenocopied *codY*::tn in butyrate but grew poorly in propionate and was not rescued by isoleucine. *lpdA1*::tn, *lpdA2*::tn, and *pdhB*::tn each grew poorly in propionate, and isoleucine supplementation likewise failed to rescue their growth. Growth of these three mutants in butyrate was higher than in propionate but still considerably worse than JE2. *lpdA1*::tn exhibited isoleucine-enhanced growth in butyrate (though less than wildtype), while *lpdA2*::tn and *pdhB*::tn did not, showing that LpdA2 or another pathway may provide redundancy to *lpdA1*::tn when butyrate is present. Together, these data support our interpretation that BCFA metabolism is disrupted by propionate and butyrate. However, the paradoxical role of IlvE in mitigating SCFA stress (i.e., it is required for optimal growth in propionate, yet its absence promotes growth in butyrate) suggests an isoleucine-independent mechanism of growth promotion in butyrate.

### Propionate and butyrate disrupt *S. aureus* membrane potential

We hypothesized that if propionate and butyrate impair *S. aureus* growth through reduced BCFA abundance, then membrane integrity would be compromised. To test this, we performed LIVE/DEAD staining on *S. aureus* grown in propionate or butyrate (the “dead” stain, propidium iodide, cannot permeate cells with a strong chemiosmotic potential across the membrane)(**Fig. 3a**). Growth in LB resulted in a SYTO9 (“live”)/PI fluorescence ratio of ∼50 arbitrary units (AU), while this ratio for cells incubated in isopropanol (control for no membrane potential) was reduced to ∼2 AU. Growth in propionate and butyrate gave ratios of ∼30 AU, indicating compromised membrane potential. To determine SCFAs altered cellular ultrastructure, we also used electron microscopy to evaluate morphological differences in the cell wall and membrane. Contrary to a recent study that reported increased peptidoglycan thickness linked to altered BCFA abundance(*31*), no obvious ultrastructural differences were observed (**Fig. 3b**).

**Fig. 3.**
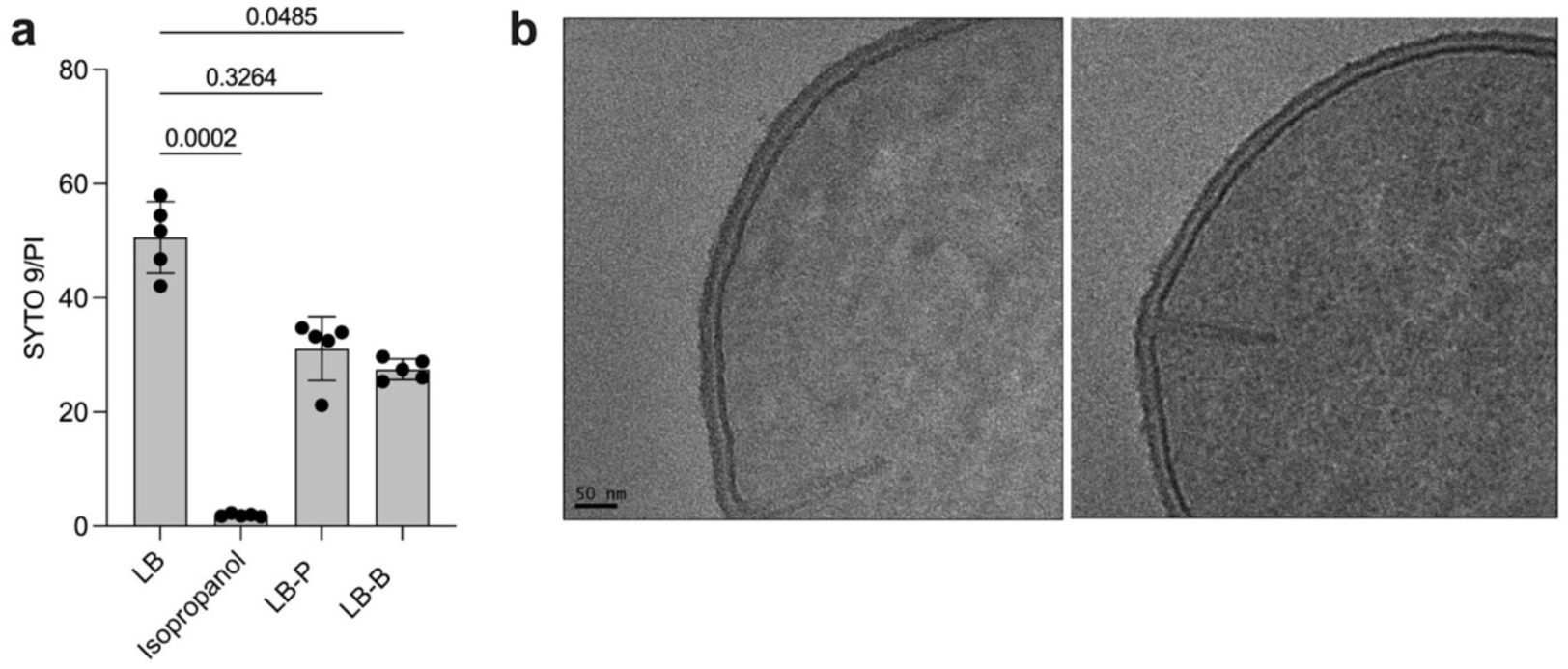
Propionate and butyrate disrupt *S. aureus* membrane potential. **A)** Ratio of fluorescence from SYTO 9 relative to propidium iodide in LIVE/DEAD stains of *S. aureus* JE2 grown in LB with or without 100 mM of sodium propionate or sodium butyrate (n=5 per condition). Isopropanol killed cells were included as a negative control. Statistical significance was determined with a Kruskal-Wallis one-way ANOVA with Dunn’s multiple comparison’s test. **B)** Representative electron micrographs of *S. aureus* JE2 grown in LB (left) or LB and propionate (right). Bar= 50nm. There were no obvious differences between growth in LB alone or LB and butyrate (image not shown).

### SCFAs invert the branched- to straight-chain fatty acid ratio in the *S. aureus* membrane

Given the compromised membrane integrity of *S. aureus* in propionate and butyrate, we performed targeted lipidomics to determine the membrane composition of JE2 when grown in LB with either SCFA, with or without exogenous BCAAs (**Fig. 4**). LB-grown *ilvE*::tn (sensitive to propionate), *lpdA1*::tn, and *lpdA2*::tn (sensitive to both SCFAs) were also evaluated. As predicted, growth of wildtype in both SCFAs resulted in a markedly decreased BCFA:straight chain FA ratio relative to LB alone **(Fig. 4a, Suppl. Fig.1)**, though ratios of specific isomer forms (branched-branched, straight/branched, and straight-straight) varied between media (**Fig. 4b,c**). For example, cells grown in butyrate exhibited decreased phosphatidylglycerol (PG) head groups with two branched acyl chains (B/B) and increased abundances of two straight acyl chains (S/S) in the 34:0, 32:0, and 30:0 isomers **(Fig. 4b)**. Growth in propionate resulted in similar reductions in B/B isomers but a large increase in S/B isomers and an increase in shorter chain S/S isomers. Consistent with growth enhancement data (**Fig. 2**), isoleucine supplementation yielded increased B/B isomers, while leucine and valine supplementation led to increased S/S isomers, although chain lengths differed between media (**Fig.4d**). Interestingly, the propionate sensitive *ilvE*::tn and *lpdA2*::tn mutants had branched/straight chain FA ratios similar to wildtype in LB alone, though the isomer composition for *ilvE*::tn was distinct (**Fig.4e**). Isomer composition in *lpdA2*::tn was comparable to wildtype, suggesting that its SCFA sensitivity may be unrelated to BCFA metabolism. *ilvE*::tn had increased PG 34:0, 32:0, and 30:0 S/B isomers, while *lpdA1*::tn exhibited lower abundances of B/B isomers and increased S/B isomers relative to wildtype and the other mutants. These data demonstrate that propionate and butyrate disrupt *S. aureus* lipid membrane homeostasis by altering BCFA metabolism, likely contributing to altered membrane integrity (**Fig.3a**).

**Figure 4.**
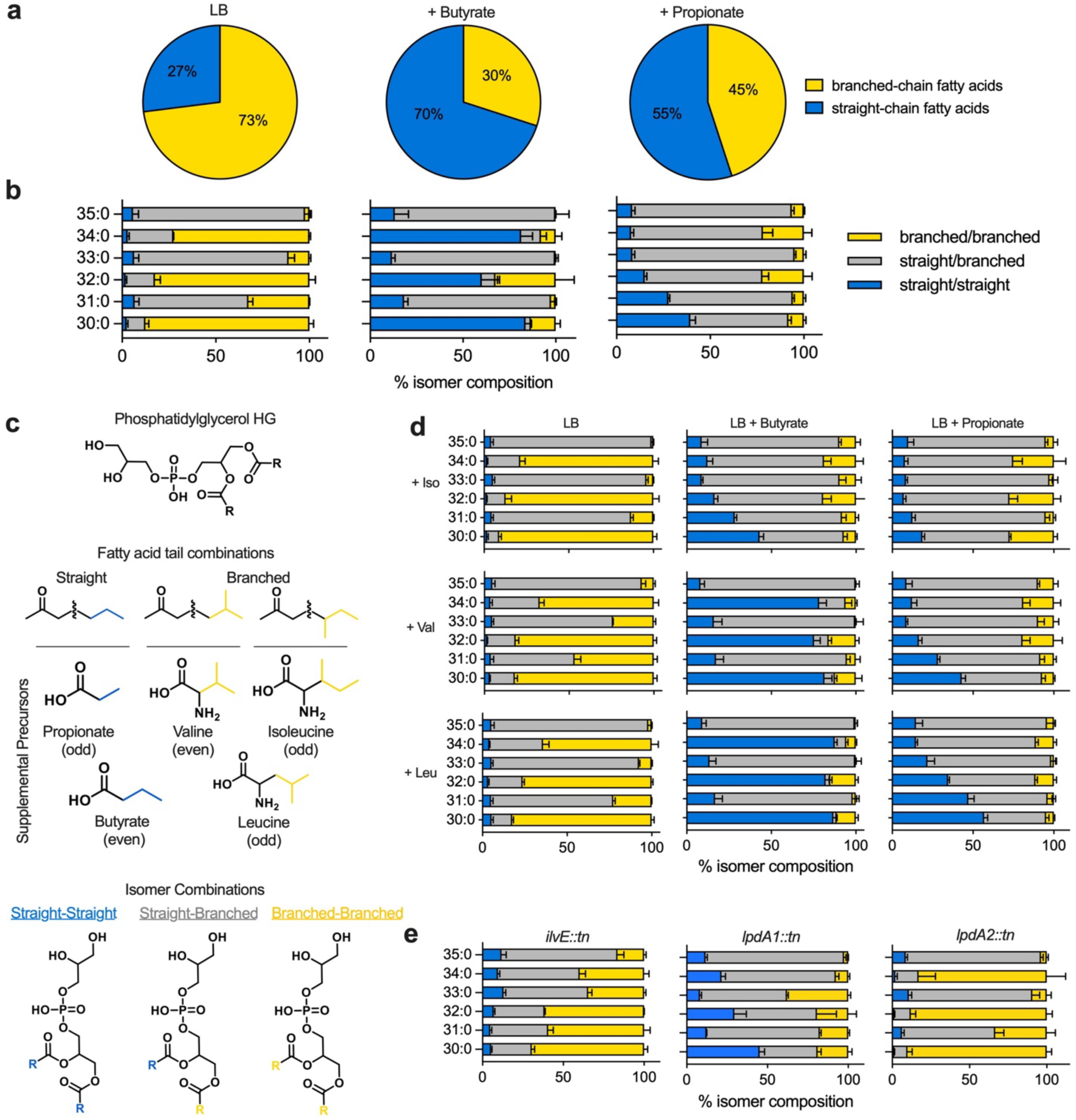
SCFAs alter *S. aureus* membrane lipid composition. **(a)** Ratio of branched-chain fatty acids to straight-chain fatty acids in *S. aureus* JE2 grown in LB with and without supplementation of sodium propionate or sodium butyrate. **(b)** Individual lipid isomer composition in LB, or LB with propionate/butyrate supplementation. For each PG (30:0-35:0) three isomers (branched-branched, branched-straight, and straight-straight fatty acid combinations, shown in panel **c**) can be expected. Data are expressed as percentages of total PGs as culture densities varied. **(d)** PG isomer percentage in LB, LB+Propionate, and LB+Butyrate supplemented with branched chain amino acids isoleucine, valine, and leucine. **(e)** Isomer composition in JE2 transposon mutants (*ilvE::tn, lpdA1::tn, lpdA2::tn*) that were more sensitive to propionate and whose growth was not rescued by exogenous isoleucine.

### SCFAs potentiate membrane-targeting antimicrobials

We next hypothesized that altered lipid composition induced by SCFAs would increase *S. aureus* sensitivity to membrane-targeting antimicrobials. To test this, JE2 was grown in LB with propionate and butyrate as before, but with escalating doses of colistin, polymyxin B, daptomycin, or the antimicrobial peptide LL-37 (cathelicidin). Each targets the bacterial cell membrane with distinct modes of action and specificities (**Fig.5a)**. For example, polymyxin B and colistin perturb the membrane by displacing stabilizing cations and increasing permeability(*32*). As these bind to lipopolysaccharides, they are considered ineffective against Gram-positive bacteria. Daptomycin is a cyclic lipopeptide primarily effective against Gram-positives by inserting its lipophilic tail into the cell membrane, driving depolarization via formation of ion-conducting pores(*33*). LL-37 binds to both Gram-positive and Gram-negative membranes via electrostatic interactions and forms disordered regions in the lipid bilayer, in turn driving leakage of cell content(*34*).

**Figure 5.**
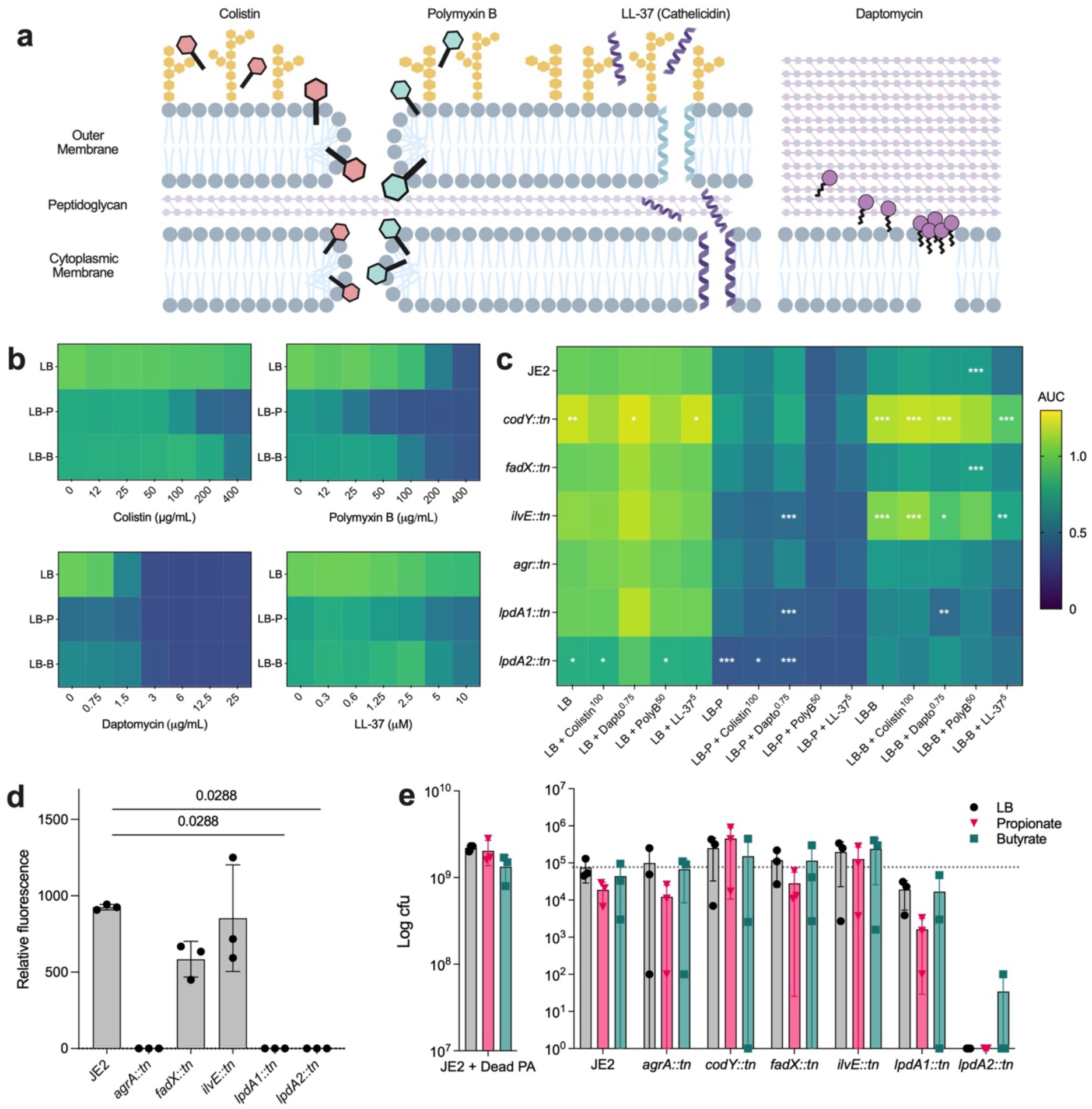
Altered BCFA metabolism by SCFAs sensitizes *S. aureus* to antimicrobial activity, impacts *agr* signaling, and reduces its fitness in competition with *P. aeruginosa*. **(a)** Graphical depiction of the proposed mechanisms of action of each antimicrobial. **(b)** Heatmaps of normalized AUCs of JE2 grown in LB with or without propionate and butyrate, supplemented with escalating concentrations of colistin (upper left), polymyxin B (upper right), daptomycin (bottom left), or the antimicrobial peptide LL-37 (bottom right). **(c)** Heatmap of normalized AUCs of JE2 and several transposon mutants in similar conditions as B, except only one dose of the antimicrobial was used. Data in B and C were normalized to JE2 in LB. **(d)** Relative fluorescence of *S. aureus* JE2 and various mutants carrying pAH1 after growth for 24 hours in LB broth. Kruskal-Wallis with Dunn’s correction for multiple comparisons was used to test for statistical significance. (e) *S. aureus* CFUs after 24 hours of co-culture with *P. aeruginosa* PA14 on permeable membranes on LB agar plates with or without 10 mM sodium propionate or sodium butyrate. Left panel depicts JE2 co-cultured with isopropanol-killed PA14. Note the difference in values on the y-axes between the left and right panels.

In LB alone, JE2 exhibited minimal sensitivity to colistin (up to 400 μg/mL) while its growth was impinged at intermediate doses of daptomycin (>1.5 μg/mL), polymyxin B (50 μg/mL) and LL-37 (5μM)(**Fig.5b)**. As predicted, JE2 exhibited greater sensitivity to each antimicrobial in the presence of SCFAs, with propionate being the more effective potentiator. Transposon mutant growth under these conditions exhibited concordance with previous experiments (**Fig.5c, Suppl. Dataset 4)**; SCFA sensitive *ilvE*::tn, *lpdA1*::tn, and *lpdA2*::tn mutants exhibited worse growth than wildtype in propionate. Consistent with their growth phenotypes (Fig. 2c), *codY*::tn and *ilvE*::tn sensitivity to colistin, daptomycin, and polymyxin B was largely unaffected, though LL-37 exhibited increased activity relative to LL-37 in LB alone. These data suggest that despite differing spectra of activity and clinical use, SCFA-induced membrane alterations may increase access of these antimicrobials to their respective targets, expose novel targets, and/or restrict antimicrobial defense mechanisms (e.g., efflux).

### Altered BCFA metabolism impairs *agr* signaling

The *S. aureus agrBDCA* operon encodes the accessory gene regulator (agr) quorum-sensing system that regulates expression of several virulence factors(*35*). We previously showed that its activity is compromised in the presence of SCFAs(*13*), leading us to hypothesize that loss of membrane homeostasis through reduced BCFAs also impairs agr signaling. Using a P3*agr-mCherry* reporter, we found that while wildtype exhibited high levels of relative fluorescence, *lpdA1*::tn signal was undetectable, similar to an *agrA*::tn control (**Fig. 5d**). Despite having a branched/straight chain FA ratio similar to wildtype (**Fig. 4e**), *lpdA2*::tn fluorescence was also undetectable, while the propionate-sensitive *fadX*::tn mutant displayed intermediate P3*agr* activity compared to JE2 and *lpdA* mutants. In contrast, and despite having significant alterations to its membrane composition, *ilvE*::tn fluorescence was similar to JE2, although it varied across replicates. These data show that BCFA homeostasis is mechanistically linked to the function of agr, and are consistent with a recent report showing diminished *agrC* expression in the *lpdA1*::tn mutant(*21*).

### *S. aureus*-*P. aeruginosa* competition is shaped by BCFA metabolism

Pathogen colonization of the airways occurs in a complex milieu of microbiota and the metabolites they exchange, mucus, and other host-derived molecules. *S. aureus* and *P. aeruginosa* coinfection of the airways is well described, as is competition between them *in vitro*, with both organisms possessing mechanisms that influence each other’s fitness(*36–38*). Here we tested the hypothesis that SCFAs tip the competitive balance in favor of *P. aeruginosa* due to compromised membrane integrity in *S. aureus*. *P. aeruginosa* PA14 and *S. aureus* (JE2 or transposon mutant) were co-cultured on permeable membranes on LB agar with or without 25mM propionate or butyrate (100 mM impairs *P. aeruginosa* growth)(**Fig. 5e**). When compared to co-culture on LB with isopropanol-killed PA14, ∼4 logs fewer *S. aureus* CFUs were recovered when live bacteria were used, indicating robust PA14-driven effects on *S. aureus* viability. Co-culture on propionate resulted in a four-fold reduction in viable *S. aureus* compared to LB, though it was not statistically significant; CFUs were also lower on butyrate but variable between replicates. *codY*::tn survived co-culture with PA14 on LB and LB + butyrate similarly to JE2 in both conditions, while higher *codY*::tn CFUs were recovered on propionate. *fadX*::tn showed no differences in viability under these conditions relative to wildtype, while *ilvE*::tn was unaffected by either SCFA. *lpdA1*::tn was more sensitive to PA14-mediated killing than JE2 in both SCFA media compared to LB alone, though neither condition was statistically significant. Finally, *lpdA2*::tn was markedly less fit in co-culture with PA14, being at or below the level of detection (10^2^ CFU) in all three media. These data show that SCFAs influence *S. aureus* competitive fitness and that BCFA metabolism contributes to its ability to resist killing by *P. aeruginosa*.

### SCFA-mediated phenotypes are conserved in CF and CRS isolates of *S. aureus*

JE2 is a derivative of USA300 LAC that was isolated from a skin abscess. Therefore, we isolated *S. aureus* from expectorated and surgically acquired airway mucus and performed assays under identical growth conditions as with JE2 (**Fig. 6**). A toxic shock syndrome isolate (MN8) was also included. We observed a variety of SCFA sensitivity phenotypes, with some strains growing similar to JE (224 and 247-01), while others were more (222, 249-01, 269-01, 337, 340, MN8) or less (255-01, 283, 296) resistant to one or both SCFAs (**Fig. 6a)**. Importantly, isoleucine supplementation enhanced growth of all strains in both SCFAs, supporting our model that propionate and butyrate disrupt *S. aureus* BCFA metabolism.

**Figure 6.**
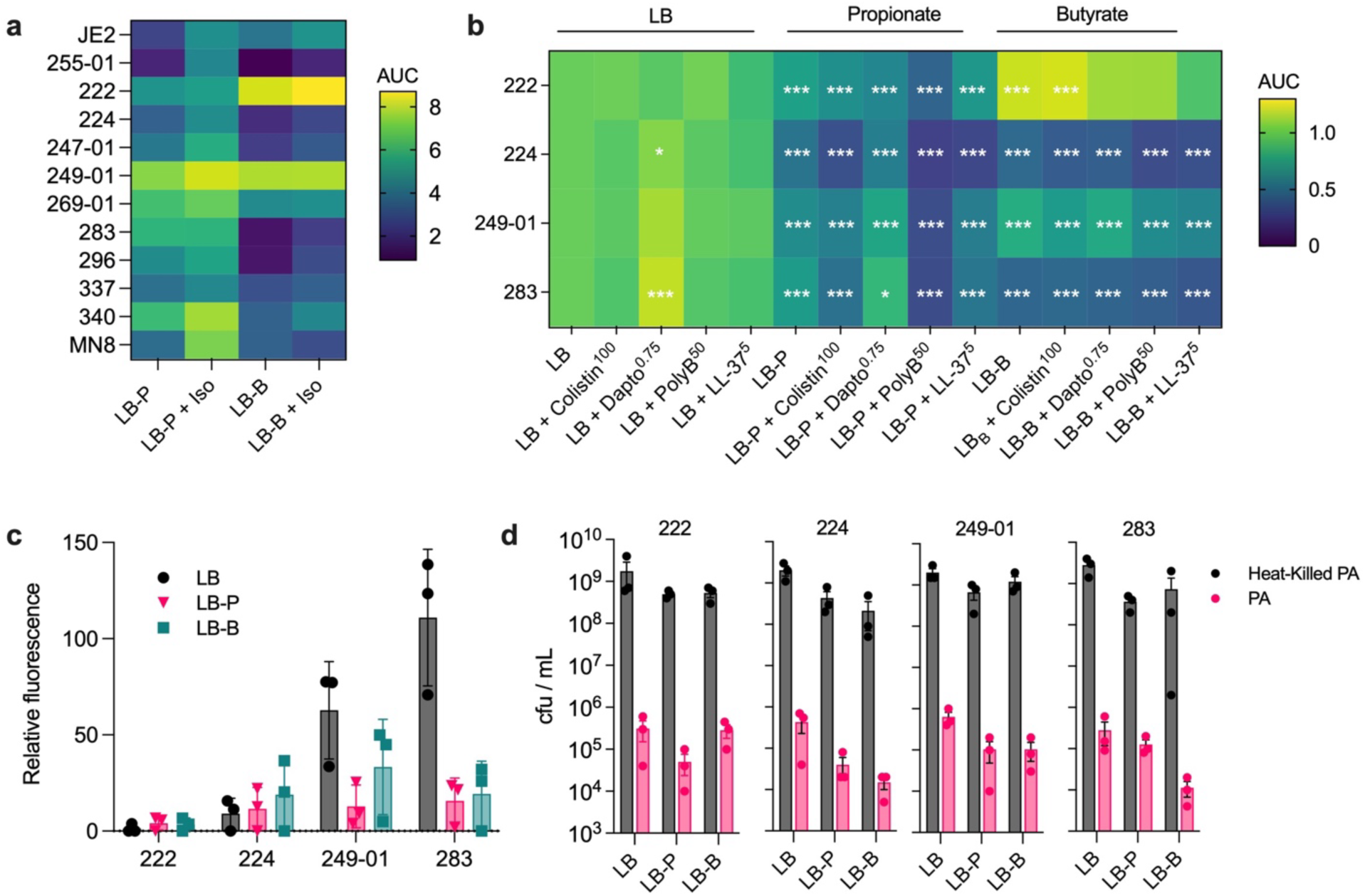
Clinical isolate data. (**a)** Ten clinical isolates of *S. aureus* from patients with CF or CRS and one from a patient with toxic shock syndrome (MN8) were grown in LB with propionate or butyrate, with and without isoleucine supplementation (n=3 biological replicates). **(b)** Antimicrobial susceptibility of clinical isolates with SCFA supplementation (n=3 biological replicates). Data in each row were normalized to the first column (untreated clinical isolate in LB). Data were compared using a two-way ANOVA with Dunnett’s multiple comparison test (*, p<0.05; **, p<0.01; ***, p<0.001) **(c)** Fluorescence of clinical isolates carrying the P3*agr-mCherry* reporter plasmid pAH1 when grown in LB with or without SCFA supplementation (n=3 biological replicates). **(d)** Fitness in competition with *P. aeruginosa* PA14 with and without the presence of SCFAs.

We narrowed our focus to strains with growth phenotypes in SCFA that were divergent from JE2 (222, 224, 249-01, 283). We performed assays on these strains carrying the P3*agr-mCherry* reporter and found that neither 222 or 224 fluoresced, suggesting nonfunctional *agr* systems (**Fig. 6c)**. This is not unexpected, as *agr*-deficiency has been reported in clinical isolates(*39, 40*). In contrast, 249-01 and 283 exhibited AgrBDCA-mediated fluorescence (though lower than JE2) which was suppressed by SCFA supplementation. Antimicrobial sensitivity in SCFAs was variable, with daptomycin having no effect on 249 in butyrate, or modestly increasing growth of 224, 249-01, and 283 in propionate (**Fig. 6b, Suppl. Dataset 5**). Strain 222 exhibited increased growth in butyrate supplemented with colistin relative to butyrate alone, although daptomycin, polymyxin B, and LL-37 led to decreased growth. The fitness of each isolate in competition with *P. aeruginosa* was also diminished by one or more SCFA **(Fig. 6d)**. Viability of 222, 224, and 249-01 were lower from co-cultures on propionate relative to LB, as were those from 224, 249-01, and 283 on butyrate. Collectively, these data support the hypothesis that SCFAs broadly impact *S. aureus* physiology by disruption of membrane homeostasis.

## Discussion

The emerging picture of airway colonization by strict and facultative anaerobes raises questions about their contributions to disease and interactions with canonical respiratory pathogens like *S. aureus*. A key class of metabolites produced by anaerobes in the airways are SCFAs, which accumulate as byproducts of carbohydrate and amino acid fermentation(*5, 10*). Here, we extend earlier findings from our group and others detailing the impaired growth of *S. aureus* in the presence of SCFAs(*11, 13, 41*). We used a genetic and multi-omic approach to show that propionate and butyrate disrupt *S. aureus* BCFA metabolism, leading to quorum signaling inhibition, increased sensitivity to membrane-targeting antimicrobials, and reduced fitness in competition with *P. aeruginosa*.

BCFAs comprise the majority of lipid species in the *S. aureus* membrane when grown in bacteriological media (e.g., LB) and the ratio of BCFAs to straight chain FAs is an essential mechanism regulating *S. aureus* membrane homeostasis and its response to environmental perturbation(*16, 17, 19, 20*). *S. aureus* BCFA mutants can bypass BCFA auxotrophy by obtaining host unsaturated fatty acids via Geh lipase activity(*42*), and presumably these host lipids would allow it to overcome growth inhibition by propionate and butyrate. However, it is unclear what role Geh plays in *S. aureus* pathogenesis in CF or CRS, as *geh* was not induced in endogenous *S. aureus* in CF sputa compared to *in vitro* conditions, nor was it induced by propionate *in vitro*(*41, 43*). Regardless, pleiotropic effects of propionate and butyrate on *S. aureus* physiology may not be solely due to altered membrane fluidity, but also the seeming importance of BCFAs to membrane protein function, as observed with the Sae two-component system(*21*). This raises the intriguing question of how *S. aureus* senses membrane composition and appropriately adjusts its metabolism to reach homeostasis in a variety of host environments.

The influence of SCFAs on membrane lipid composition also highlights BCAA availability in a host as a source of essential BCFA precursors. Kaiser et al. showed the importance of *S. aureus* BCAA uptake via BcaP and BrnQ1 for persistence in a murine nasal colonization model(*44*), although mice were pretreated to deplete endogenous microbiota. In an environment with competition between the microbiota for limited nutrients, SCFA-producing anaerobes in close physical proximity to *S. aureus* could diminish the fitness of the latter by slowing its growth and production of quorum-regulated virulence factors. Intuitively, this may be exacerbated by low levels of available BCAAs.

SCFAs may also modulate the host response, as instillation of micro-to-millimolar concentrations of propionate to the murine lung prior to challenge with luminescent *S. aureus* led to increased luminescence in treated mice relative to untreated mice after 6 hours; this was interpreted as greater *S. aureus* growth due to a blunted inflammatory response(*45*). Conversely, CF bronchial epithelial cells treated with SCFAs *in vitro* secrete more IL-8 than non-CF cells(*10*). In addition to SCFAs produced locally in the airways by anaerobes, systemically circulating SCFAs produced by the gut microbiota may also affect pathogen colonization and the host response. Indeed, propionate and butyrate protected mice against *S. aureus* in a model of experimental mastitis by strengthening the blood-milk barrier(*46*).

Our studies were performed on *S. aureus in vitro* with supraphysiological levels of SCFAs, therefore they are not direct models of microbial community interactions and evolution. Rather, they provide insights for future research into plausible mechanisms governing *S. aureus* colonization patterns over the lifetime of a patient with chronic airway disease, including its membrane biogenesis and homeostasis. We posit that SCFA biocompatibility coupled with their ability to potentiate antimicrobials make them attractive tools to study control of *S. aureus in vivo*.

## Methods

### Bacterial strains and propagation

*S. aureus* JE2, the plasmid-free derivative of USA300 LAC, and transposon insertion mutants from the JE2 Nebraska Transposon Mutant Library(*47*) were routinely propagated in LB broth (IBI Scientific IB49030) and on LB agar (Fisher BioReagents BP1423) at 37°C, with shaking at 220 rpm. Erythromycin (4 μg/mL) and chloramphenicol (10 μg/mL) were added as necessary to select for transposon mutants or strains carrying pAH1, respectively. Strains used are shown in Table S1. Clinical isolates of *S. aureus* were obtained by streaking clinically derived sinus mucus onto mannitol salt agar (MSA) and incubating aerobically overnight at 37°C.

### Patient recruitment and mucus collection

Patients with a positive diagnosis of CRS undergoing functional endoscopic sinus surgery (FESS) were recruited at the University of Minnesota Clinics and Surgery Center. Sinus secretions were obtained from a single maxillary sinus under endoscopic visualization by suction into Argyle mucus traps (Cardinal Health, Dublin, OH). Individuals with cystic fibrosis were also recruited during routine outpatient visits at the University of Minnesota Cystic Fibrosis Center during which they provided a single expectorated sputum sample. Samples were processed for *S. aureus* isolation as described above. Protocols were approved by the UMN Institutional Review Board (1403M49021 and 1404M49426).

### Growth curves with SCFAs, BCAAs, or antibiotics

LB-grown plate and liquid cultures of *S. aureus* JE2 and transposon mutants were diluted 1:100 in PBS, from which 5 μL was added to 195 μL of experimental media in a flat-bottom 96-well plate. Media were LB, LB supplemented with 100 mM of either sodium propionate (Sigma P1880) or sodium butyrate (Sigma 303410), LB supplemented with 1 mg/mL of one BCAA (isoleucine, leucine, or valine), or LB supplemented with either SCFA and one of the three BCAAs. Additional growth assays were performed in SCFA LB with and without varying concentrations of colistin sulfate, polymyxin B, daptomycin, or cathelicidin (LL-37). For the growth curves with daptomycin, 50 μg/mL of calcium chloride was added to both LB or LB plus daptomycin. 96-well plates were then incubated in a BioTek Synergy H1 microplate reader for 24 hours at 37°C. Plates were subjected to 5 seconds of orbital shaking prior to hourly readings at OD600. The area under the curve (AUC) for each biological replicate was calculated in Graphpad Prism with the following settings: Y = 0, peaks less than 10% of the distance from minimum to maximum Y were ignored, and all peaks must go above the baseline of 0.

### LIVE/DEAD staining

*S. aureus* was cultured overnight in LB and diluted either 1:500 into fresh LB or 1:50 in LB containing SCFAs. 1:50 was used to account for the SCFA-induced growth impediment. Cultures were incubated for approximately 4 hours at 37°C with shaking at 220 rpm. Per the LIVE/DEAD BacLight (Invitrogen L7012) protocol, cultures were centrifuged at 10,000 rpm for 10 minutes. Pellets were then washed 3X with 0.85% NaCl and allowed to incubate on the benchtop for 1 hour. Next, cells were stained with SYTO-9/propidium iodide (3 μL of dye mix per 1 mL of cells in 0.85% NaCl) and allowed to incubate in the dark at room temperature for 15 minutes. 100 μL of each sample was added to a black-walled, flat-bottom 96-well plate in triplicate. Fluorescence was determined using a BioTek Synergy H1 plate reader with an excitation wavelength of 485 nm and emission wavelengths of 530 nm (green) and 630 nm (red). Data are reported as the ratio of green fluorescence to red fluorescence.

### Transmission electron microscopy

*S. aureus* JE2 was grown in LB, LB + propionate, or LB + butryate for 6h, washed three times with 50mM HEPES buffer, enrobed in 2% Noble agar, cut into 1-2mm blocks and chemically fixed with 2% glutaraldehyde in HEPES buffer for 2h. Cells were washed thrice prior to additional fixation and stained en bloc using 2% (wt/vol) osmium tetroxide in HEPES buffer for 2h, followed by staining with 1% (wt/vol) uranyl acetate for 1h. Samples then underwent serial dehydration in 25%, 50%, 75%, 95% ethanol for 15 minutes each, followed by three additional incubations in 100% anhydrous ethanol. Agar blocks were then suspended in a 50:50 solution of a 100% ethanol:LR White resin solution for ∼2 hours, followed by 100% LR White for an additional 2 hours. Blocks were embedded in gelatin capsules containing fresh LR White and were allowed to polymerize at 60°C for 2 hours. Blocks were thin-sectioned on a Reichert-Jung Ultracut E microtome and mounted on formvar and carbon-coated 200 mesh copper grids. To improve contrast, sections were post-stained in 1% uranyl acetate, prior to imaging on a FEI Tecnai G2 F20 transmission electron microscope at the Advanced Analysis Center at the University of Guelph (Canada).

### RNA extraction

Overnight LB cultures of *S. aureus* were diluted 1:500 into fresh media and grown until the OD600 reached ∼0.2-0.3. Cells were pelleted by centrifugation at 14,000 rpm for one minute, then resuspended in 50 μL of LB supplemented with 20 ug/mL of lysostaphin (Sigma-Aldrich L7386) and incubated at 37°C for 15 minutes. Lysates were dissolved in 1 mL of TRIzol Reagent (ThermoFisher) and incubated on the benchtop for 5 minutes, followed by addition of 200 μL of chloroform (VWR). Samples were agitated by hand for 15 seconds, allowed to sit on the benchtop for 5 minutes, then centrifuged at 12,000 rpm for 15 minutes at 4°C. The aqueous phase (∼500-550 μL) was removed and mixed with an equal volume of 95% ethanol and vortexed for 5 seconds. The mixture was then subjected to on-column DNase I treatment and clean-up using the Zymo RNA Clean & Concentrator-5 kit, following the manufacturer’s protocol. RNA concentration was determined using the Qubit Broad Range RNA kit (ThermoFisher).

### NanoString quantification of gene expression

RNA extracted from *S. aureus* under various growth conditions was submitted to the University of Minnesota Genomics Center, where it was hybridized to probes from a custom codeset targeting transcripts for various virulence factors, transcriptional regulators, and metabolism, as well as six housekeeping transcripts<u>21</u>. Differential expression analysis was performed using the NanoString nSolver Advanced software. Transcripts were considered differentially expressed if their Benjamini-Hochberg adjusted p-value was less than 0.05.

### Lipid extraction and analysis

Membrane lipids were extracted using a modified Bligh and Dyer method, as described previously(*31, 48*). Briefly, *S. aureus* pellets were suspended in LC-MS grade water at McFarland standards that differed by 0.04 units or less. A 2 mL portion of each suspension was transferred to a glass centrifuge tube, sonicated on ice for 30 min, and 2mL of a chilled 2:1 methanol/chloroform solution was added. After 5 min of periodic vortexing, 0.5 mL of chilled chloroform and 0.5 mL of chilled water were added, followed by vortexing for 1 min and centrifugation at 4°C and 2,000 rpm for 10 min. The lipid-containing organic layer was collected into new glass tubes, dried in a speed-vac, reconstituted with 0.5 mL of 1:1 chloroform/methanol. Samples were stored at -80°C in sealed glass vials. For lipid analysis, 40μL of each extract was transferred to a new vial, dried in a speed-vac, and reconstituted with 100μL of mobile phase A (MPA, see below) for negative mode (2.5X dilution) or 40μL for positive mode (1X dilution).

Lipid separations were performed on a Waters Acquity FTN I-Class Plus ultra-performance liquid chromatography (UPLC) system equipped with a Waters Acquity charged surface hybrid (CSH) C18 (2.1 x 100mm, 1.7um) column, as described previously(*49*). Mobile phase A (MPA) consisted of 60:40 acetonitrile/water with 10 mM ammonium formate. Mobile B phase (MPB) consisted of 88:10:2 isopropanol/acetonitrile/water with 10 mM ammonium formate. LysylPGs were analyzed separately using mobile phase solutions adjusted to pH 9.7 with ammonium hydroxide. All solvents and salts were LC-MS grade. A 30 min gradient elution was performed with a flow rate of 0.3mL/min using the following conditions: 0-3 min, 30% MPB; 3-5 min, 30-50% MPB; 5-15 min, 50-90% MPB; 15-16 min, 90-99% MPB; 16-20 min, 99% MPB; 20-22 min, 99-30%; 22-30 min 30% MPB. Column temperature was maintained at 40°C and samples were kept at 6°C. Injection volume was 10μL.

The UPLC was connected to the electrospray ionization (ESI) source of a Waters Synapt XS traveling wave ion mobility mass spectrometer (TWIM-MS). The following parameters were used for ESI: Capillary voltage, +/- 2.0; sampling cone voltage, 40V; source offset, 4V; source temperature, 120°C; desolvation temperature, 450°C; desolvation gas flow, 700L/hr; cone gas flow, 50L/hr. Traveling wave separations were performed using a wave velocity of 550 m/s, wave height of 40V, and a nitrogen gas flow of 90mL/min. The time-of-flight (TOF) mass analyzer was operated in V-mode (resolution mode) with a resolution of ∼30,000. Mass calibration was performed with sodium formate over a mass range of 50-1200 m/z. Data were collected over the 30 min separation with 0.5s scan time. Data-independent acquisition (DIA) of MS/MS (MSE) was performed in the transfer region of the instrument using a 45-60 eV collision energy ramp. Leucine enkephalin was continuously infused throughout for post-acquisition mass correction.

Lipidomic data were analyzed using the small molecules version of Skyline software(*50*). Transition lists containing fatty acid tail composition of 30:0-35:0 were created for PGs (negative mode, [M-H]-adducts) and LysylPGs (positive mode, [M+H] adducts). Each lipid species in the transition list contained FA fragments ranging from 13 to 20 carbons. Waters .raw files were imported directly into Skyline. Lock mass correction was performed using a 0.05 Da (negative mode) or 0.1 Da (positive mode) window. Ion mobility filtering was performed with a drift time width of 0.60 ms. Chromatograms from MS1 and MS/MS dimensions were overlaid and used to identify fatty acyl tails of lipid species based on matching retention and drift times. Peak areas for each lipid species, isomer, and fatty acid fragment were integrated to evaluate differences in *S. aureus* membrane composition between growth conditions.

### Fluorescent *agr* reporter activity

P3*agr-mCherry* reporter activity was assayed as described previously(*13*). Briefly, *S. aureus* JE2, transposon mutants, and clinical isolates of *S. aureus* were electroporated with pAH1 and plated on LB agar with 10 μg/mL chloramphenicol (cam^10^) to select for transformants. Isolated colonies were streaked onto fresh LB cam^10^ agar prior to use in experiments. Cultures were grown at 37°C for 24 hours with shaking at 220 rpm. 200 μL from each culture was added to a black-walled, flat-bottom 96-well plate in technical duplicates. Fluorescence was measured on a BioTek Synergy H1 microplate reader using excitation and emission wavelengths of 580 nm and 635 nm, respectively. Fluorescence was divided by the OD600 for each well, and the technical replicates were averaged to yield the relative fluorescence for each biological replicate (n=3).

### Bacterial competition assays

Competition assays between *P. aeruginosa* and *S. aureus* were performed using established protocols(*51*). *S. aureus* JE2, a series of JE2 transposon mutants, *S. aureus* clinical isolates, and *P. aeruginosa* PA14 were grown overnight in LB broth at 37°C with shaking at 220 rpm. JE2 transposon mutant cultures were maintained under selection with 4 μg/mL of erythromycin. After overnight growth, each strain was sub-cultured 1:5 in pre-warmed fresh LB and allowed to grow for another ∼2 hours. The OD600 was determined and cultures were centrifuged at 5,000 rpm for 5 minutes in an Eppendorf benchtop 5418 centrifuge. Supernatants were discarded and each pellet was suspended in 1 mL of sterile PBS, then diluted in PBS to OD600 = 0.01 (*S. aureus*) and OD600 = 0.02 (*P. aeruginosa*), resulting in an approximate density of 107 CFU/mL of each bacterium. These were serially diluted and plated on LB agar for enumeration on the following day. In parallel, one culture of PA14 was centrifuged as described, then suspended in 1 mL of isopropanol for ∼30 minutes. It was then washed three times in sterile PBS to create a dead PA control. For the competition assay, 50 μL of diluted PA14 was added to 50 μL of diluted *S. aureus* (or a transposon mutant or clinical isolate) in a new 0.5 mL Eppendorf tube and the mixture was vortexed for 5 seconds. 5 μL of the mixture was spotted onto a Nucleopore Track-Etch membrane (0.2 μm pore size, 13 mm diameter) that had been placed onto the surface of an LB agar plate using sterile tweezers. Plates were incubated agar-side up at 37°C for 24 hours, after which the membranes were removed with sterilized tweezers and suspended in 1 mL of PBS. These were vortexed for 5-10 seconds at maximum speed, then allowed to sit on the benchtop for ∼10 minutes, followed by another 5-10 seconds of vortex and manual disruption of any remaining consolidated biomass via pipetting. Once uniformly dispersed, samples were serially diluted in PBS and plated onto mannitol salt agar (MSA) and Pseudomonas isolation agar. These plates were incubated overnight at 37°C and colonies were enumerated the following day.

## Acknowledgements

We thank Todd Markowski and LeeAnn Higgins at the UMN Center for Mass Spectrometry and Proteomics, and Cezar Khursigara and Erin Anderson at the University of Guelph Advanced Analysis Center for their assistance with TEM. We also acknowledge Dr. Holly Boyer, Erin Feddema, Anika Tella, and Ali Stockness in the Department of Otolaryngology at UMN, and the staff at the UMN Cystic Fibrosis Center for their assistance with patient recruitment, IRB development, and sample collection. This work was supported by a National Institute for Allergy and Infectious Disease (NIAID) Research Project Grant (5R01AI177613-03) and Cystic Fibrosis Foundation Research Grant (HUNTER22P0) awarded to RCH. JRF was supported by a CF Foundation postdoctoral fellowship (FLETCH21F0), RM was supported by a NHLBI T32 Fellowship (2T32HL007741-21), and ARV was supported by an Administrative Research Supplement (HL136919-03S1). CDF and KMH were supported by NIAID Research Project Grant (R01AI173144).

