## Supplemental Information for "Commensal-derived short-chain fatty acids disrupt lipid membrane homeostasis in *Staphylococcus aureus*"

Supplemental Material

**Table S1.** Strains used throughout this study.

| ***Staphylococcus aureus*** | | | | | | | |
| --- | --- | --- | --- | --- | --- | --- | --- |
| **Strain** | **Locus tag** | **NTML ID^1^** | | | **BEI ID** | | **Source^2^** |
| JE2 |  |  | | | NR-46543 | | 1 |
| *agrA::tn* | SAUSA300_1992 | NE1532 | | | NR-48074 | | 1 |
| *bcaP::tn* | SAUSA300_2538 | NE206 | | | NR-46749 | | 1 |
| *brnQ1::tn* | SAUSA300_0188 | NE945 | | | NR-47488 | | 1 |
| *brnQ2::tn* | SAUSA300_0306 | NE605 | | | NR-47148 | | 1 |
| *brnQ3::tn* | SAUSA300_1300 | NE44 | | | NR-46587 | | 1 |
| *codY::tn* | SAUSA300_1148 | NE1555 | | | NR-48097 | | 1 |
| *crtM::tn* | SAUSA300_2499 | NE1444 | | | NR-47986 | | 1 |
| *fadX::tn* | SAUSA300_0229 | NE263 | | | NR-46806 | | 1 |
| *fakA::tn* | SAUSA300_1119 | NE229 | | | NR-46772 | | 1 |
| *fakB1::tn* | SAUSA300_0733 | NE1540 | | | NR-48082 | | 1 |
| *fakB2::tn* | SAUSA300_1318 | NE403 | | | NR-46946 | | 1 |
| *ilvD::tn* | SAUSA300_2006 | NE718 | | | NR-47261 | | 1 |
| *ilvE::tn* | SAUSA300_0539 | NE292 | | | NR-46835 | | 1 |
| *leuB::tn* | SAUSA300_2011 | NE76 | | | NR-46619 | | 1 |
| *lpdA1::tn* | SAUSA300_1467 | NE1896 | | | NR-48438 | | 1 |
| *lpdA2::tn* | SAUSA300_0996 | NE1610 | | | NR-48152 | | 1 |
| *pdhB::tn* | SAUSA300_0994 | NE1758 | | | NR-48300 | | 1 |
|  | **Description** | |  |  | |  | |
| MN8 | Methicillin sensitive isolate from a patient with toxic shock syndrome | | | | | | 3 |
| 222 | Clinical isolate from patient with chronic sinusitis | | | | | | 4 |
| 224 | Clinical isolate from patient with chronic sinusitis | | | | | | This study |
| 247-01 | Clinical isolate from patient with chronic sinusitis | | | | | | This study |
| 249-01 | Clinical isolate from patient with chronic sinusitis | | | | | | This study |
| 255-01 | Clinical isolate from patient with chronic sinusitis | | | | | | This study |
| 269-01 | Clinical isolate from patient with chronic sinusitis | | | | | | This study |
| 283 | Clinical isolate from patient with chronic sinusitis and cystic fibrosis | | | | | | This study |
| 296 | Clinical isolate from patient with chronic sinusitis | | | | | | This study |
| 337 | Clinical isolate from patient with chronic sinusitis | | | | | | This study |
| 340 | Clinical isolate from patient with chronic sinusitis | | | | | | This study |
| ***Pseudomonas aeruginosa*** | | | | | | | |
| PA14 | UCBPP-PA14, burn wound isolate | | | | | | 5 |

^1^ Nebraska Transposon Mutant Library mutant ID

^2^ JE2 and transposon mutants were provided by the Network on Antimicrobial Resistance in *Staphylococcus aureus* (NARSA) for distribution through BEI resources, NIAID, NIH: *Staphylococcus aureus* subsp. *aureus* Strain JE2, NR-46543

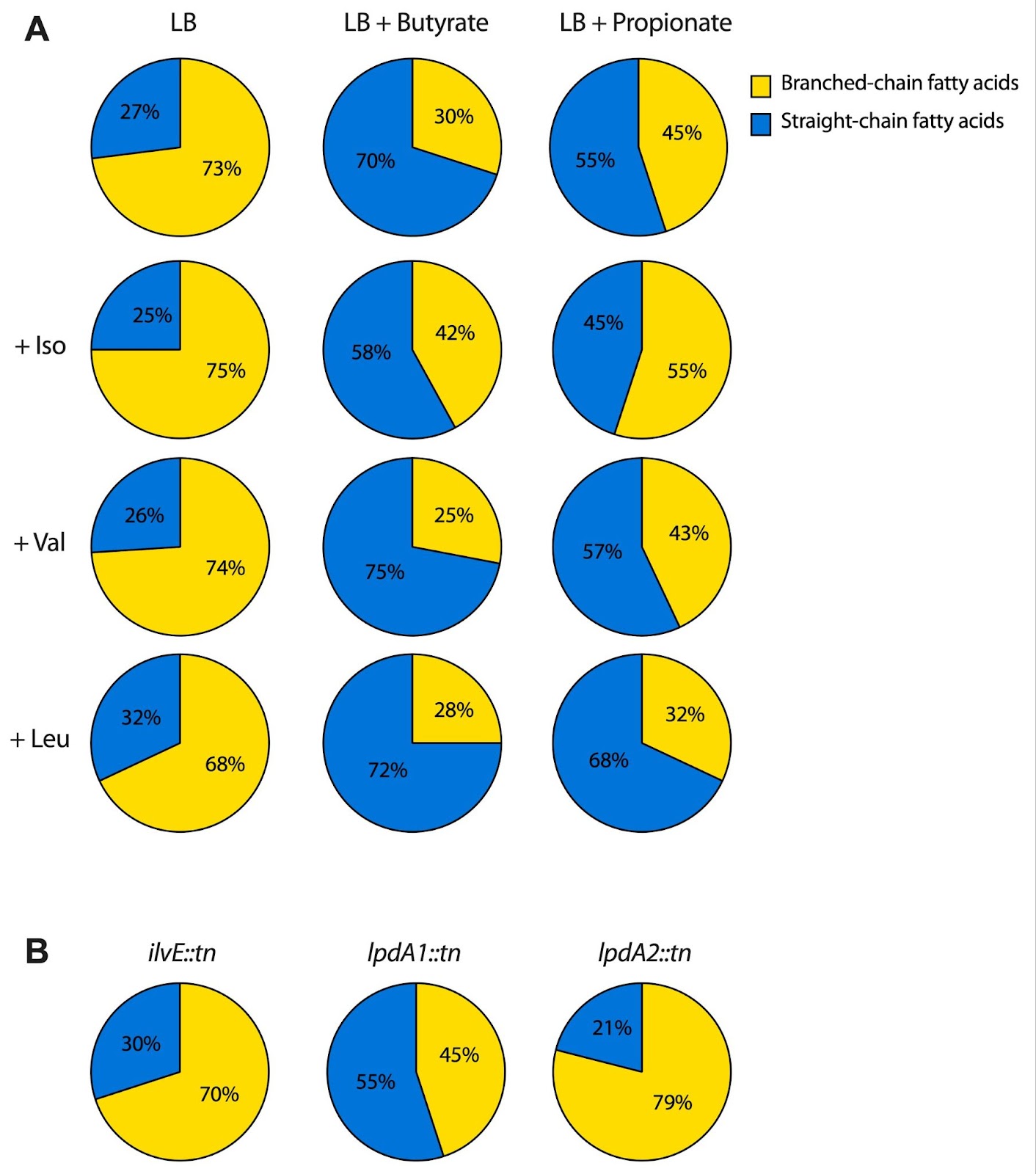

**Figure S1. SCFAs alter *S. aureus* membrane lipid composition. (A)** Ratio of branched-chain fatty acids to straight-chain fatty acids in *S. aureus* JE2 grown in LB with and without supplementation of sodium propionate or sodium butyrate, with or without supplementation of 1mg/mL BCAAs isoleucine (iso), valine (val), or leucine (leu). **(B)** ratio of branched-chain fatty acids to straight-chain fatty acids in *S. aureus* JE2 transposon mutants from the Nebraska Transposon Mutant Library/
